# SurfaceGenie: a web-based application for prioritizing cell-type specific marker candidates

**DOI:** 10.1101/575969

**Authors:** Matthew Waas, Shana T. Snarrenberg, Jack Littrell, Rachel A. Jones Lipinski, Polly A. Hansen, John A. Corbett, Rebekah L. Gundry

## Abstract

**Motivation:** Cell-type specific surface proteins can be exploited as valuable markers for a range of applications including immunophenotyping live cells, targeted drug delivery, and in vivo imaging. Despite their utility and relevance, the unique combination of molecules present at the cell surface are not yet described for most cell types. A significant challenge in analyzing ‘omic’ discovery datasets is the selection of candidate markers that are most applicable for downstream applications.

**Results:** Here, we developed *GenieScore*, a prioritization metric that integrates a consensus-based prediction of cell surface localization with user-input data to rank-order candidate cell-type specific surface markers. In this report, we demonstrate the utility of *GenieScore* for analyzing human and rodent data from proteomic and transcriptomic experiments in the areas of cancer, stem cell, and islet biology. We also demonstrate that permutations of *GenieScore*, termed IsoGenieScore and OmniGenieScore, can efficiently prioritize co-expressed and intracellular cell-type specific markers, respectively.

**Availability and Implementation:** Calculation of *GenieScores* and lookup of *SPC scores* is made freely accessible via the SurfaceGenie web-application: www.cellsurfer.net/surfacegenie.

## Introduction

Cell surface proteins play key roles in diverse biological processes and disease pathogenesis through mediation of adhesion and signaling between the extracellular and intracellular space. Owing to their accessible location, cell surface proteins can be exploited as valuable markers for a range of research and clinical applications including immunophenotyping live cells, targeted drug delivery, and *in vivo* imaging. As such, a growing interest in cell-type specific data has fueled the generation of the Cell Surface Protein Atlas (Bausch-Fluck et al., 2015), Human Protein Atlas (Uhlen et al., 2005), Human Cell Atlas Project (Regev et al., 2017), and related efforts. However, the unique combination of molecules present specifically at the cell surface are not yet described for most cell types or disease states, and thus continued innovation regarding surface protein discovery and annotation efforts are needed.

Specialized chemoproteomic approaches which specifically label and enrich cell surface proteins can provide direct evidence of cell surface localization resulting in empirically-derived snapshot views of the cell surface proteome (Kalxdorf, Gade, Eberl, & Bantscheff, 2017; Turtoi et al., 2011; Wollscheid et al., 2009). However, the large sample requirements and technical sophistication required for these experiments preclude their widespread use and application to sample-limited cell types. For these reasons, whole-cell proteomic and transcriptomic-based approaches that can be applied to identify and quantify thousands of molecules from fewer cells will continue to be useful in the search for cell surface proteins that are informative for particular cell types or disease stages.

Independent of the discovery strategy employed, bioinformatic predictions can serve as an important complement to experimental approaches by providing a means to filter data and prioritize the focus on proteins that are predicted to be localized to the cell surface (Bausch-Fluck et al., 2018; da Cunha et al., 2009; Diaz-Ramos, Engel, & Bastos, 2011; Town et al., 2016). Though transcriptomic and proteomic approaches offer significant advantages over antibody screening with regards to throughput and depth of coverage, candidate markers identified from discovery approaches must be subsequently validated as viable immunodetection or payload-delivery targets. Considering the significant cost and time required for the development of *de novo* affinity reagents, it is prudent to select candidates in a manner that considers whether a marker is likely to be both accessible and detectable by affinity reagents in a manner that allows cell types of interest (*i.e.* target cells) to be discriminated from non-target cells. Moreover, these assessments should be objective and suited to the analysis of large datasets such as those generated by proteomic and transcriptomic studies. To address these outstanding needs, we developed *GenieScore*, a metric to rank and prioritize candidate cell type specific surface markers - calculated by integrating a consensus-based prediction of cell surface localization with user-input quantitative data. Here, we describe the development of *GenieScore* and demonstrate its utility for prioritizing candidate cell surface markers using data obtained from proteomic workflows that specifically identify cell surface proteins (*e.g.* Cell Surface Capture (CSC)) and more general strategies (*e.g.* whole-cell lysate proteomics and transcriptomics). We also demonstrate that permutations of *GenieScore*, termed IsoGenieScore and OmniGenieScore, can efficiently prioritize co-expressed and intracellular cell-type specific markers, respectively. To facilitate its implementation among users, we developed SurfaceGenie, an easy-to-use web application that calculates *GenieScores* for user-input data and annotates the data with ontology information particularly relevant for cell surface proteins. SurfaceGenie is freely available at www.cellsurfer.net/surfacegenie.

## Results

### Generation of a surface prediction consensus dataset for predictive localization

Four previous bioinformatic-based constructions of the human cell surface proteome were compiled into a single, surface prediction consensus (SPC) dataset containing 5,407 protein accession numbers (Dataset S1.1). The strategies used to generate these predicted human surface protein datasets varied markedly, from manual curation to machine learning, and resulted in dataset sizes ranging from 1090 to 4393 surface proteins (Figure 1A). Despite these differences, there was considerable overlap among these predictions, with 69% and 41% of proteins in the SPC dataset occurring in ≥ 2 or ≥ 3 individual prediction sets, respectively. The number of proteins exclusive to a prediction strategy is positively correlated to the original dataset size, albeit not linearly, comprising 1.7%, 4.4%, 9.6%, and 26.5% for the Diaz-Ramos, Bausch-Fluck, Town, and Cunha datasets, respectively (Figure 1B). To reflect the difference in the consensus of surface localization, each protein was assigned one point for each of the individual predicted datasets in which that protein appeared, termed *Surface Prediction Consensus* (*SPC) score* (Figure 1B, Dataset S1.1). The distribution of *SPC score*s is shown in Figure 1B where 1671, 1507, 1497, and 732 proteins are assigned a score of 1, 2, 3, and 4, respectively. To enable more widespread applicability, mouse and rat *SPC score* databases were generated by mapping the human proteins to mouse and rat homologs using the Mouse Genome Informatics database (http://www.informatics.jax.org, Dataset S1.2-3).

**Figure 1:**
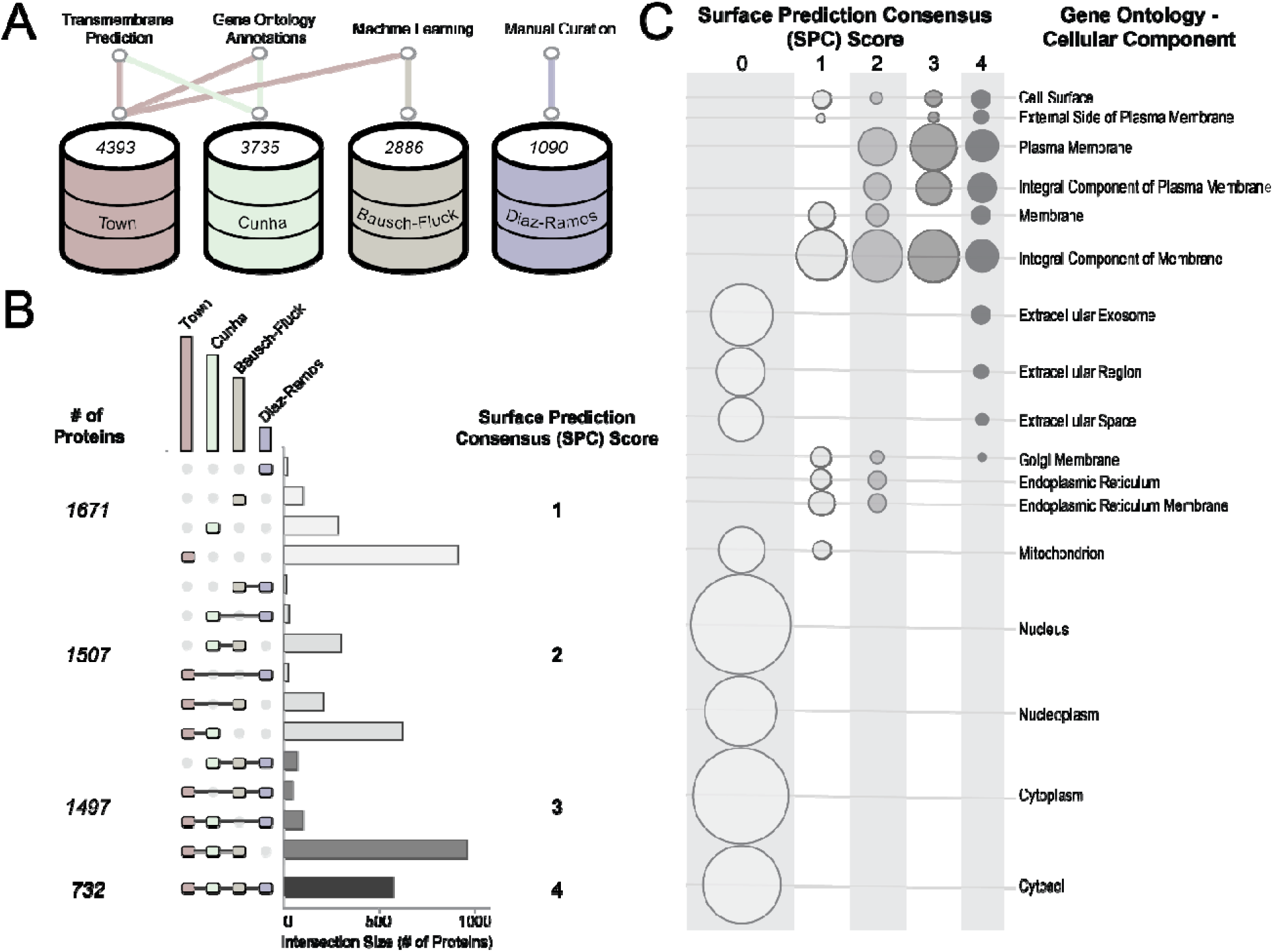
Generation and benchmarking of a *Surface Prediction Consensus* (*SPC*) score. (A) The four previously published human surfaceome databases used, designated by first author of the corresponding publication, with details about how the databases were generated and the number of UniProt Accessions within each database. (B) An UpSet plot (Conway, Lex, & Gehlenborg, 2017; Lex, Gehlenborg, Strobelt, Vuillemot, & Pfister, 2014) depicting the intersections between the individual surfaceome databases. The proteins were stratified by the number of individual datasets they appeared in, termed *Surface Prediction Consensus* (*SPC*). The number of proteins with each SPC score is shown. The full dataset is provided in the Supporting Information (Dataset S1, 4.1) (C) The distribution of Gene Ontology Cellular Component Ontology (GO-CCO) annotations across different *SPC scores* depicted as a bubble chart, where the size of the bubble represents the number of proteins in the intersection between the particular *SPC score* and GO-CCO annotations.

### Benchmarking the SPC dataset against other annotations

The SPC dataset was compared to three established resources for obtaining cell surface localization annotations: 1) Gene Ontology Cellular Component Ontology (GO-CCO) (Ashburner et al., 2000; The Gene Ontology, 2019), 2) annotations within the Cell Surface Protein Atlas (CSPA) (Bausch-Fluck et al., 2015), and 3) annotations generated through application of HyperLOPIT (Christoforou et al., 2016). Comparisons to GO-CCO were consistent with expectations as ‘nucleus’ and ‘cytoplasm’ were the two most common terms for proteins with *SPC scores* of 0, ‘integral component of membrane’ and ‘membrane’ for *SPC scores* of 1, and ‘integral component of membrane’ and ‘plasma membrane’ for *SPC score*s of 2-4 (Figure 1C). The ‘confidence’ assignment to proteins in the CSPA agreed with *SPC scores* for both human and mouse, with the notable exception of ∼17% of proteins assigned ‘high confidence’ having an *SPC score* of 0 (Figure S1A). However, upon closer inspection, 95% of these proteins are predicted to be secreted or extracellular matrix proteins (Secretome P, (Bendtsen, Kiemer, Fausboll, & Brunak, 2005)), which can be captured in CSC experiments but are not integral membrane proteins. The most common HyperLOPIT annotation in proteins with *SPC scores* of 3 or 4 was ‘plasma membrane’; however, ‘ER/Golgi apparatus’ was the most common annotation in proteins with *SPC scores* of 1 or 2 (Figure S1B). Though these comparisons demonstrated agreement overall, the SPC dataset provides unique and specific information in addition to assigning the predictions in a non-binary manner. Furthermore, as the *SPC score* is not dependent on experimental observation, it is more comprehensive in coverage than the CSPA and HyperLOPIT. These differences offer significant advantages for mathematically assigning the likelihood that a protein is present at the cell surface in a predictive manner. Moreover, the calculation of *SPC score* is straightforward and flexible to allow easy integration of results from future efforts of cell surface localization prediction.

### Defining features of a cell surface protein marker based on first principles

By defining the term ‘marker’ to designate a cell surface protein which is capable of distinguishing between cell types of interest on the basis of signal obtained by immunodetection, there are three features that can be used to evaluate the capacity of a protein to serve as a marker (Figure 2A). These include (1) *SPC score* - presence at the cell surface, (2) *signal dispersion* - difference in abundance among cell types, and (3) *signal strength* - scaling factor to account for the aim of obtaining specific antibody-based detection. The product of these three terms, which we define as *GenieScore*, is a metric that can be used to rank proteins from experimental data for their capacity to serve as a marker. Importantly, prioritization of cell surface proteins that are likely capable of serving as informative markers should consider experimental data from relevant cell types, including the target and non-target cell types that are to be discriminated. Hence, although a consensus-based predictive approach can be adopted to represent whether a protein is capable of being present at the cell surface (*SPC score*), the *signal dispersion* and *signal strength* must be determined empirically, as these will differ among cell types. Additional descriptive details and rationalization for the *GenieScore* components are provided in Supporting Information along with examples of *GenieScore* calculations for different experimental observations (Figure S2B). As the two mathematical terms chosen to represent *signal dispersion* and *signal strength* are agnostic to the data type, we investigated the effects of different sources of input-data to these terms with respect to the calculated *GenieScores*.

**Figure 2.**
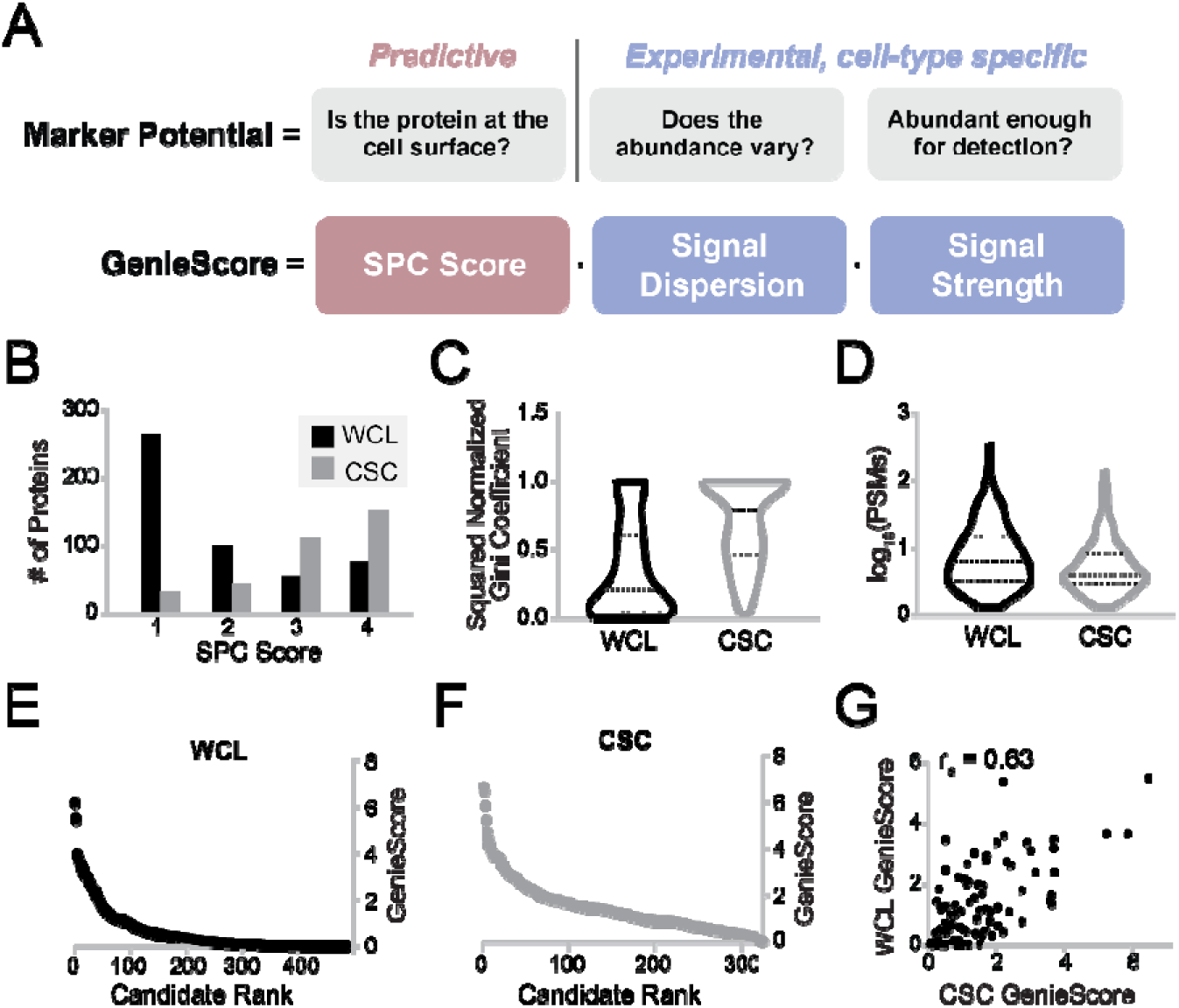
*GenieScore* components and application to two proteomic analyses of fou lymphocyte lines. (A) The features of a protein that were hypothesized to predominate the capacity of a protein to serve as a cell surface marker are shown with the names of the mathematical terms derived to represent them. The marker potential features are annotated by the applied approach (i.e. predictive or experimental) to answer the relevant questions. The remaining panels depict the distribution of the individual components and *GenieScores* calculated from the data acquired from application of whole-cell lysate (WCL) or Cell Surface Capture (CSC) to four lymphocyte cell line (*n* = 3 per cell line, N = 485 data points for WCL, N = 325 data points for CSC). (B) A histogram depicting the distribution of *SPC scores* within predicted surface proteins (SPC score >0) identified by application of WCL and CSC. (C) A violin plot depicting the distribution of *signal dispersion* for the predicted surface proteins identified by WCL and CSC. (D) A violin plot depicting the distribution of *signal strength* for the predicted surface proteins identified by WCL and CSC. (E) Plot of *GenieScore* against rank-order of candidate cell surface markers for predicted surface proteins identified by WCL. (F) Plot of *GenieScore* against rank-order of candidate cell surface markers for predicted surface proteins identified by CSC. (G) *GenieScores* calculated using either WCL or CSC data are plotted against each other the 91 proteins identified by both approaches along with the Spearman’s Correlation for those scores.

### Testing GenieScore by comparing two proteomic approaches for marker discovery

We previously demonstrated that CSC applied to four human lymphocyte cell lines resulted in sufficient depth of surface protein identification to allow discrimination among the lines (Haverland et al., 2017). Here, we performed whole-cell lysate (WCL) digestion of these same cell lines to determine whether a generic proteomic approach would be sufficient to detect divergent cell surface proteins to do the same. Notably, the WCL approach only required a peptide amount equivalent to ∼1000 cells compared to CSC which used ∼100 million cells/experiment. While the majority, 75% (325), of the CSC-identified proteins are predicted to be cell surface localized (*i.e. SPC score*s of 1-4), only 13% (485) of the WCL proteins met this criterion. Although these datasets were collected on the same cell lines, only 91 proteins with *SPC score*s 1-4 were observed in both datasets, which represent 28% and 19% of the CSC and WCL predicted surface proteins, respectively. Despite these differences, applying hierarchical clustering to the subset of proteins in each dataset with *SPC scores* of 1-4 recapitulated the clustering predicted based on the entire dataset for both proteomic approaches (Figure S3). These data highlight the utility of applying the SPC metric as a strategy to filter and compare between datasets and demonstrate that a generic proteomic strategy can provide sufficient predicted surface protein detection to differentiate among cell types using 0.1% of the cellular material required for CSC, albeit without the empirical evidence of localization for the detected set predicted surface proteins provided by application of CSC.

*GenieScores* were calculated for each protein in the CSC and WCL datasets using peptide-spectrum matches (PSMs) as input for *signal dispersion* and *signal strength* calculations (Dataset S.1-2). Though predicted surface proteins were identified by both proteomic approaches, the distributions of *SPC scores, signal dispersion,* and *signal strength* were markedly different between CSC and WCL (Figure 2B-D). These observations are expected due to the highly-selective nature of CSC which primarily captures *N-*glycosylated peptides resulting in higher specificity for *bona fide* surface proteins and fewer peptides identified per protein (Boheler et al., 2014; Gundry et al., 2012; Haverland et al., 2017; Wollscheid et al., 2009). *GenieScores* were plotted against the rank for CSC and WCL data resulting in a rectangular-hyberbola-like shape, namely a subset of higher-scoring proteins that trail off into a majority of lower-scoring proteins (Figure 2E-F) with a similar range (6.59 and 6.16 for CSC and WCL, respectively) but significant difference in the distribution. *GenieScores* for the 91 proteins identified in both proteomic approaches were strongly correlated (r_s_ = 0.63) (Dataset S2.3, Figure 2G).

The top-scoring candidate markers in both the CSC and WCL data sets are proteins for which the majority (if not the totality) of PSMs are in a single cell line. The numbers of PSMs per cell line for selected proteins are shown for CSC and WCL in Figure 3 along with the ranks determined by application of *GenieScore* (plotted in Figure 2E-F). Many of the high ranking candidates have previously been reported as markers for cancer types modeled by the cell lines used here - including ATP1B1, CD39, and HLA-DR for chronic lymphocytic leukemia (CLL) (Damle et al., 2002; Johnston et al., 2018; Pulte et al., 2007) (Hg-3 cell line); CD10 and CD79b for Burkitt Lymphoma (He et al., 2018; “WHO Classification: Tumours of the Haematopoietic and Lymphoid Tissues (2008),” 2016) (Ramos cell line) (Figure 3). Proteins with moderate ranks often had PSMs spread evenly among two or more of the cell lines. The examples here include CD5 - a known T cell marker that is often expressed in CLL (Damle et al., 2002) (accounting for its observation in Jurkat and Hg-3 cell lines, respectively), and CD47 – a protein reported to be upregulated in many cancer subtypes (Takimoto et al., 2019). Proteins with low ranks are equally spread among all the cell lines, often ‘housekeeping’ type proteins such as transferrin receptor 1 and mannose-6-phosphate receptor (Figure 3).

**Figure 3.**
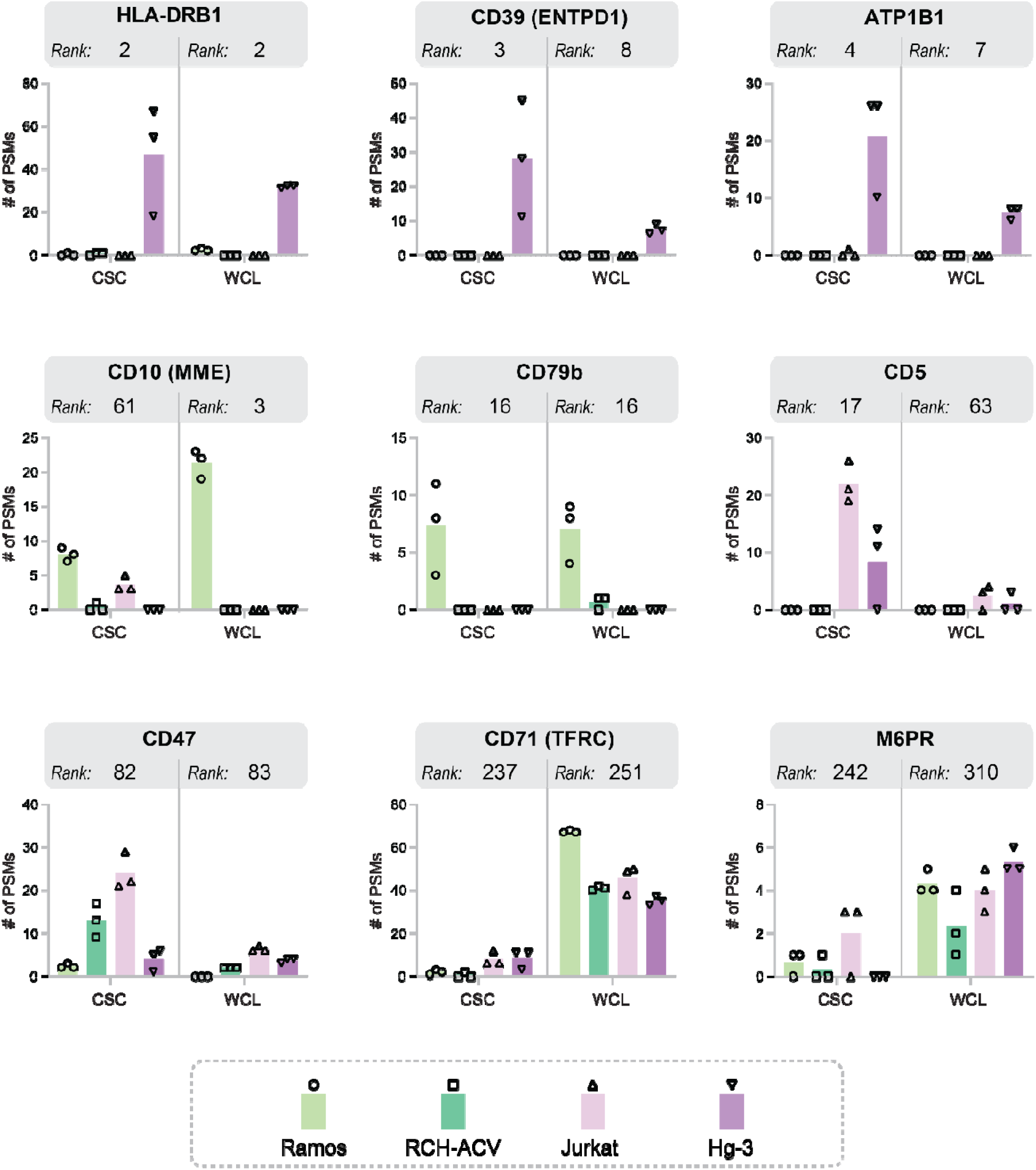
Distributions of observed abundance for selected proteins in the lymphocyte data with a range of *GenieScores*. The number of peptide-spectrum matches (PSMs) assigned to selected proteins for both Cell Surface Capture (CSC) and whole-cell lysate (WCL) experiments. Biological replicates (*n* = 3) are shown as data points and averages are shown as columns. The ranks assigned to each protein, according to the set of calculated *GenieScores*, are shown for both CSC and WCL datasets.

As the calculation of *GenieScore* relies on averages (as opposed to individual replicate measurements) the relationship between the product of the experimental terms (*signal dispersion* and *signal strength*) used to calculate *GenieScore* and the statistical difference (which considers variability in measurement) between cell lines was investigated. A positive relationship was observed, with Spearman’s correlations (r_s_) of 0.66 and 0.64 for WCL and CSC, respectively, suggesting that the equation for *GenieScore* is likely to be prioritizing proteins for which there is a statistical difference (Figure S4A). Finally, recognizing the limitations of relying on PSMs for quantitative comparisons, *GenieScores* were calculated using MS1 peak areas for selected proteins. Results from this strategy had a very strong correlation to *GenieScores* using PSMs; r_s_ = 0.88 and 0.80 for WCL and CSC, respectively (Figure S4B). Altogether, application of *GenieScore* to the data collected from diverse lymphocyte cell lines produced similar ranking of candidate markers independent of the type of input data, including several surface proteins previously linked to relevant cancer subtypes, which indicates *GenieScore* is a robust and valid prioritization metric.

### Benchmarking GenieScore against two published surface protein marker studies

A major application of *GenieScore* is to prioritize candidate markers for immunophenotyping. Hence, we sought to benchmark the performance of *GenieScore* ranking against two published studies that performed flow cytometry analyses to orthogonally validate putative markers for cell types of interest which were originally identified from proteomics and/or transcriptomic data. In the first study, Martinko *et al.* performed CSC and RNA-Seq on MCF10A KRAS^G12V^ cells (comparing the results to empty vector control MCF10A cells) to identify surface proteins indicative of RAS-driven cancer phenotype (Martinko et al., 2018). Antibodies were subsequently developed against seven candidate markers, all of which demonstrated positive signal on the MCF10A KRAS^G12V^ cells. Using *GenieScore* as a prioritization metric, we investigated the relative ranks of these validated markers among the CSC and RNA-Seq datasets. As the goal of the original study was to identify surface proteins which were upregulated in cancer, *GenieScores* were only calculated for predicted surface proteins (*SPC score* >0) which met this additional criterion – resulting in 122 candidates from CSC and 330 candidates from RNA-Seq (Figure 4A, Dataset S2.1-2). The validated proteins were among the highest scoring candidates in both the CSC and RNA-Seq data sets (Figure 4A). *GenieScores* calculated from the CSC and RNA-Seq data had a moderate correlation (r_s_ = 0.41) with most of the validated markers scoring highly for both CSC and RNA-Seq (Figure 4B). The rank of the putative markers by *GenieScore* was compared to the rank by log_2_ fold changes (a metric denoted as selection criteria in the original manuscript) (Figure 4C). In all but one case, the candidates rank higher by *GenieScore* than by log_2_fold ratio. These results support *GenieScore* as a useful, single metric that enables selection of cell surface proteins which can serve as markers for immunodetection. Though the validated markers were among the top-ranking candidates, other proteins with high *GenieScores* emerge as potential targets, highlighting the utility of *GenieScore* to reveal new biological insights or targets from previously published data. Two such targets that scored well by both CSC and RNA-Seq are THBD and NRP1, which have been previously implicated in a KRAS-driven myeloid malignancy and KRAS-driven tumorigenesis (Meyerson et al., 2017; Vivekanandhan et al., 2017). Additionally, several proteins score well in one dataset (*i.e.* CSC or RNA-Seq) but are completely absent from the other. SAT-1, ranked 5 within the RNA-Seq data but not observed by CSC, plays a role in a polyamine synthesis pathway that is upregulated by KRAS-driven cancers (Arruabarrena-Aristorena, Zabala-Letona, & Carracedo, 2018; Linsalata et al., 2004). Inspection of the primary sequence of SAT-1 reveals that the *N*-glycosite is located within a peptide that would make it unlikely to be detected by mass spectrometry. Conversely, there are proteins within the CSC dataset for which there were no matching transcripts such as LRP1, a protein associated with tumorigenesis and tumor progression (Chen et al., 2010; Xing et al., 2016). Altogether, these results present the complementary nature of CSC and RNA-Seq as discovery techniques and demonstrate that *GenieScore*-based analyses of these data, either independently or together, provide a rapid strategy for prioritizing candidates for immunophenotyping.

**Figure 4.**
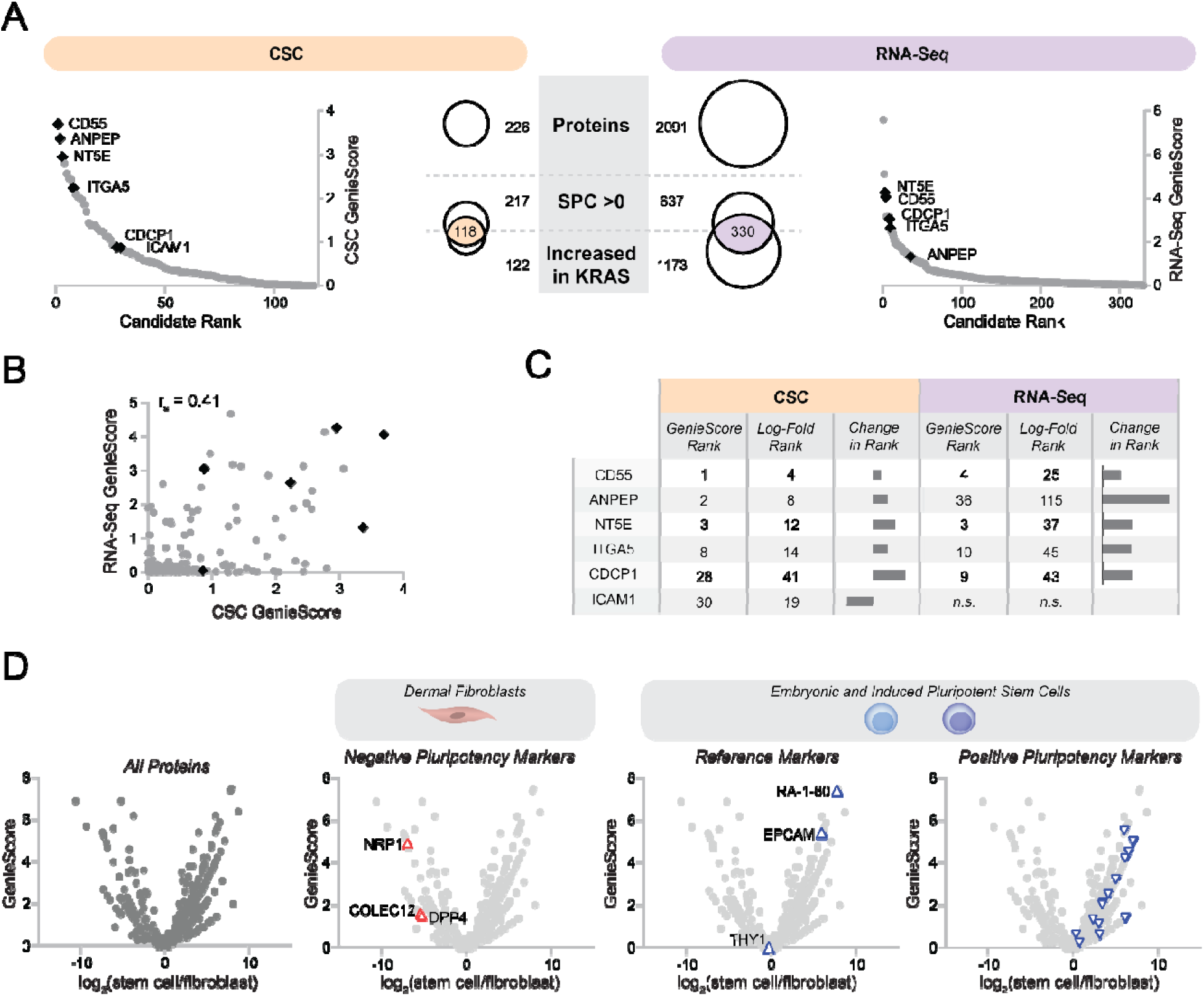
Benchmarking *GenieScore* against two published cell surface marker studies which validated candidate markers by flow cytometry. Panels A-C depict data from application of *GenieScore* to data from Martinko *et al*. Panel D depicts data from application of *GenieScore* to data from Boheler *et al.* (A) The subset of proteins for which *GenieScores* were calculated is the intersection of the set of proteins with SPC scores >0 with the set of proteins that were increased in the KRAS mutant - shown by the shaded overlap. Plots of *GenieScores* against candidate rank are shown for the Cell Surface Capture (CSC) and RNA-Seq datasets. Proteins selected in the original manuscript for antibody development and subsequently validated as surface markers by flow cytometry are shown as black diamonds and labeled with gene names. (B) The *GenieScores* calculated using either CSC or RNA-Seq data are plotted against each other for the 211 surface proteins identified by both approaches along with the Spearman’s Correlation of those scores. The flow cytometry-validated markers are shown as black diamonds. (C) A table containing the ranks assigned, according to either *GenieScores* or log_2_fold, for each protein. The change in rank, calculated as *GenieScore* rank minus log_2_fold rank, is shown for each flow cytometry-validated marker. (D) *GenieScores* for the 495 proteins identified by CSC in human fibroblast and stem cells are plotted against the log_2_fold ratio. Refence stem cell markers, as well as the negative and positive markers for pluripotency selected for validation by flow cytometry are highlighted in their own plots.

In the second study, Boheler *et al*. performed CSC on human fibroblasts, embryonic stem cells, and induced pluripotent stem cells to identify surface markers for stem cells (Boheler et al., 2014). Candidate pluripotency markers were selected by comparing the set of proteins observed on stem cells to CSC data from the CSPA, specifically, requiring that a protein was not detected in fibroblasts and detected in fewer than four other somatic cell types (excluding cancer cell types). Negative markers of pluripotency were selected in a similar manner, specifically, not detected in stem cells and detected in six or more non-diseased cell types in the CSPA. Flow cytometry analysis of human fibroblasts and stem cells was used to orthogonally validate seventeen putative positive and three putative negative pluripotency markers and included three previously reported positive pluripotency markers as controls. The *GenieScores* for proteins observed in the human fibroblast and stem cell CSC experiments were plotted against the log_2_fold ratio of PSMs between the cell types providing a visual depiction of capacity to serve as a marker segregated by cell type wherein the reference, putative negative, and putative positive pluripotency markers are denoted in individual plots (Figure 4D, Dataset S3.1). The reference stem cell markers are among the top scoring (with ranks of 2 and 10) candidate pluripotency markers from the CSC dataset, except for Thy1, a protein for which both CSC and flow cytometry results provided evidence for its presence in fibroblasts and stem cells. The *GenieScores* for the putative negative and positive markers were spread over a greater range in this dataset compared to the distribution observed for the Martinko *et al.* study. This difference is likely attributable to the notably divergent strategies employed for candidate selection. Specifically, the Boheler study relied on qualitative (presence/absence) rather than quantitative comparisons, considered data from cell types outside those included in the study, and restricted validation to candidates for which commercially-available monoclonal antibodies were available. Notably, several of the validated markers were identified by relatively few PSMs in the original dataset (IL27RA, EFNA3). While the number of PSMs is sometimes used as a filter to eliminate proteins from consideration, in this case, comparisons to 50 other cell types suggested these candidates are putatively restricted to stem cells. Thus, despite being identified by relatively few PSMs in CSC analyses, proteins that are uniquely observed in a single cell type can be valuable immunophenotyping markers provided there are data of a similar type and quality on other cell types for comparison. Altogether, these data highlight the importance of context during marker selection and the value of considering additional datasets. Specifically, if additional datasets are integrated prior to calculation of *GenieScore,* candidates with a lower *signal strength* (few PSMs) would rank more highly because they would have a higher *signal dispersion* (all PSMs coming from a single cell type). Overall, these evaluations of previously validated datasets illustrate how *GenieScore* is a useful strategy to prioritize candidate cell surface markers using both proteomic and transcriptomic datasets.

### Integrating GenieScores of proteomic and transcriptomic data to reveal candidate markers for mouse islet cell types

As *GenieScore* provided a useful rank-ordering of potential protein markers from both RNA-Seq and CSC data that was consistent with published results, we sought to evaluate its utility for integrating data from disparate studies for marker discovery. To this end, we performed CSC on mouse α and β cell lines and compared the results to published RNA-Seq data acquired on primary α and β cells from dissociated mouse islets (Benner et al., 2014). The datasets shared 321 predicted surface proteins (Figure 5A, Dataset S4.1), and *GenieScores* from the CSC data were plotted against *GenieScores* from the RNA-Seq data (Figure 5B). A possible explanation for the weak correlation (r_s_ = 0.26) between *GenieScores* is that the CSC dataset was acquired on cell lines and the RNA-Seq dataset was acquired on primary cells. However, in the context of marker discovery, each of these approaches offers advantages, namely, the CSC data provides experimental evidence regarding protein abundance at the cell surface and the RNA-Seq analysis of primary cells avoids possible artifacts introduced by culturing cells *ex vivo*. Recognizing the complementary benefits of these approaches, the data were combined in a manner that weights them equally, namely, the *signal dispersion* was calculated using the average of the normalized CSC and normalized RNA-Seq data. The combined *GenieScores* were distributed similarly to scores calculated using CSC or RNA-Seq individually and when plotted against the log_2_fold ratio between α and β cells allow for visual discrimination of the candidate markers for each cell type (Figure 5C, Dataset S4.2). Among the top candidate markers for α and β cells revealed by this combined approach are proteins with well-established roles in islet biology including GLP1, GABBR2, GALR1, KCNK3, SLC7A2 -proteins highlighted in a recent review of the β cell literature (Rorsman & Ashcroft, 2018). ALCAM (CD166), CHR1, and CEACAM1 are proteins which have been studied in the context of the islet biology, though have less defined roles (DeAngelis et al., 2008; Fujiwara et al., 2014; Schmid et al., 2011). Altogether, *GenieScore* provided a useful framework for integrating proteomic and transcriptomic data for surface marker prioritization.

**Figure 5.**
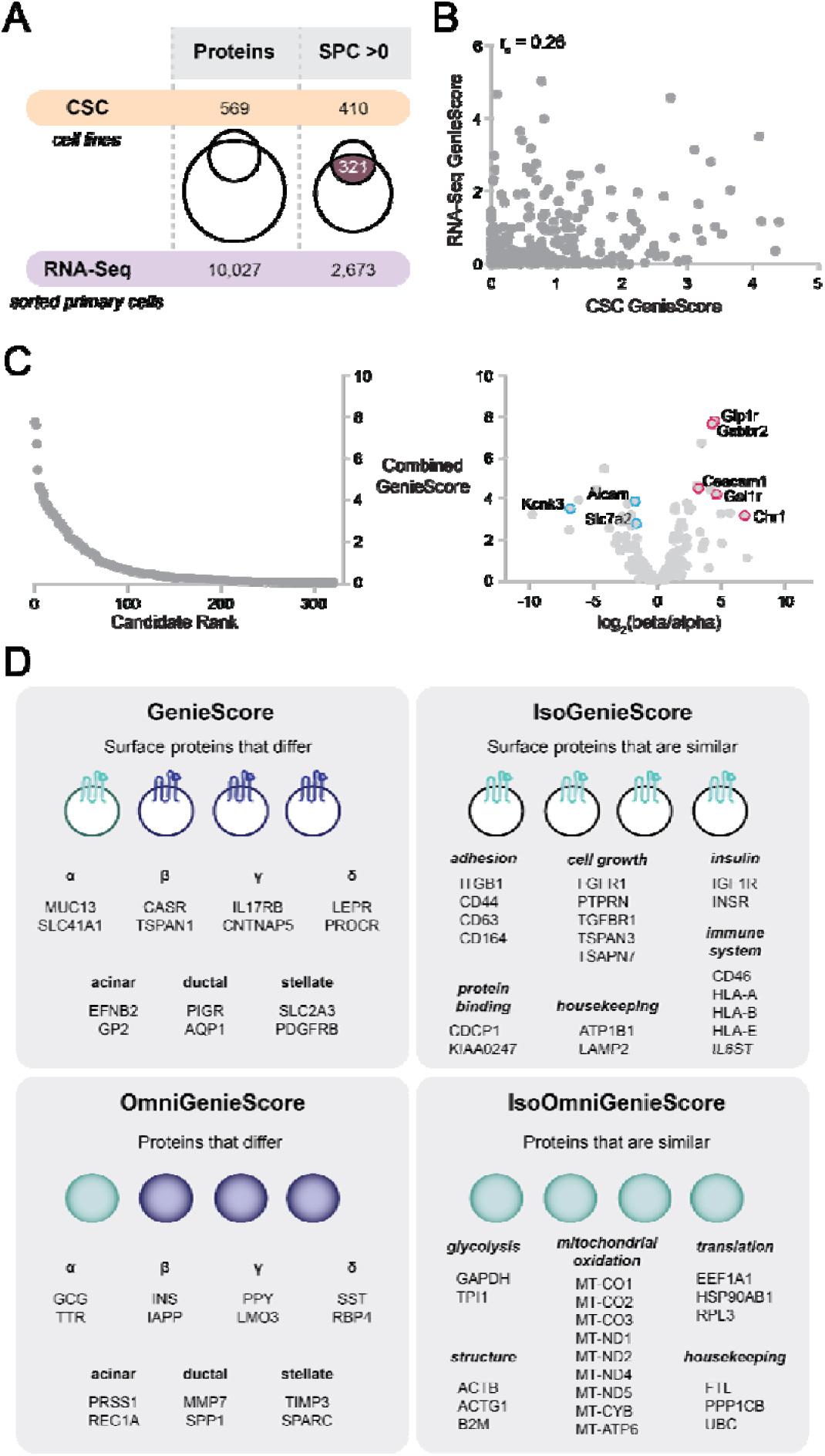
Application of *GenieScore* and its permutations to islet cell types. Panels A-C depict data from application of *GenieScore* to Cell Surface Capture (CSC) data from mouse α and β cell lines collected as part of this study integrated with RNA-Seq on mouse primary α and β cells from Benner *et al*. Panel D depicts data from application of *GenieScore* and its permutations to human islet single-cell RNA-Seq data from Lawlor *et al.* (A) The subset of proteins for which *GenieScores* were calculated is the set of proteins with *SPC scores* >0 that were identified by both CSC and RNA-Seq, shown as the shaded overlap. (B) The *GenieScores* calculated using either CSC or RNA-Seq data are plotted against each other for the 321 proteins identified by both approaches along with the Spearman’s Correlation of those scores. (C) *GenieScores* calculated using the combined CSC and RNA-Seq data are plotted against candidate rank and against the log_2_fold ratio (N = 321 proteins). Selected candidate markers which have previously been associated with islet cell biology are labeled with gene names. (D) The top scoring proteins from application of the different permutations of *GenieScore* are shown grouped either by cell type or by biological function.

To extend the analysis beyond the identification of proteins which might be capable of distinguishing α and β cells to finding cell-type specific markers within the context of the islet, we applied *GenieScore* to a single-cell RNA-Seq dataset that was collected on cells from dissociated human islets (Lawlor et al., 2017) (Dataset S5.1). Lawlor *et al* partitioned the data on single cells into seven different cell types – α, β, γ, δ, acinar, ductal, stellate - based on a subset of genes that were determined to be representative of each of the clusters. Top ranking markers for each of the seven cell types are listed in Figure 5D. Many of the proteins identified as capable of distinguishing between α and β cells in the analysis of CSC and RNA-Seq data were not cell-type specific when data from other cell types found within the islet were considered. For example, NRCAM and SLC4A10 are proteins more abundant in β than α cells, but the levels expressed in β cells are equivalent to γ or δ cells, respectively. PTPRK is expressed at a higher level in α than β cells in all studies, but the level of expression is 26-fold lower than acinar cells and 35-fold lower than ductal cells. Altogether, while the cell-type specificity that is ultimately required will depend on the desired downstream application, these observations highlight that consideration of a larger cellular context is important for the identification of cell-type specific markers.

Recognizing the utility of the *GenieScore* approach for prioritizing cell-type specific surface proteins, the equation was further adapted to enable prioritization of other classes of proteins using the same input data. First, removal of the *SPC score* component from *GenieScore*, a permutation termed *OmniGenieScore,* allowed for the identification of proteins which can be used as cell-type specific markers without considering their surface localization.

Application of *OmniGenieScore* to the islet cell single-cell RNA-seq data revealed many known cell-type specific markers such as glucagon (GCG) for α cells, insulin (INS) for β cells, pancreatic polypeptide (PPY) for γ cells, and somatostatin (SST) for δ cells (Figure 5D). By inverting the *signal dispersion* term (*i.e.*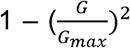), a permutation termed *IsoGenieScore*, the set of cell surface proteins which are relatively abundant and similar in signal among all cell types in the analysis were prioritized. The classes of proteins (*e.g.* adhesion, cell growth, insulin signaling) which were at the top of this ranking system were largely involved in generic processes that are not specific to any cell type (Figure 5D). Reversing the *signal dispersion* and ignoring the *SPC score*, termed *IsoOmniGenieScore*, resulted in prioritization of proteins typically selected to be loading controls for Western blotting or reference genes for PCR (*e.g.* GAPDH, ACTB, B2M) in addition to many of the proteins involved in mitochondrial oxidation (Figure 5D). Altogether, these four permutations of the *GenieScore* enabled the prioritization of candidate markers for a broad range of applications, including cell surface and intracellular markers that distinguish cell types as well as those that are co-expressed among cell types.

### SurfaceGenie: a web-based application for integrating GenieScore and relevant annotations

To enable calculation of *GenieScores* for user input data, a shinyApp, SurfaceGenie, was developed in R. In this interface, users upload data from proteomic or transcriptomic experiments as a .csv file and can view the distribution of *GenieScores* and *SPC scores* for the proteins contained in their data (Figure 6A). SurfaceGenie is compatible with human, mouse, and rat data. As part of the analysis, input proteins are annotated with ontological information including Cluster of Differentiation (CD) and Human Leukocyte Antigen (HLA) molecule designations. In addition, proteins are annotated with the number of cell types within the CSPA in which the protein has been observed – a factor found to be relevant for marker prioritization in the Boheler *et al* data. The plots and data generated are available for download, including the results for individual terms used to calculate *GenieScore*. The permutations of *GenieScore* applied in Figure 5D are also available. Additional functionality includes the ability to query accession numbers in single or batch mode, independent of data type, to obtain *SPC Scores*. SurfaceGenie is freely available at http://www.cellsurfer.net/surfacegenie (screen captures shown in Figure 6B). A User Guide with step-by-step tutorials is provided in the Supporting Information; it can be alternatively accessed through the web application or the Github repository.

**Figure 6.**
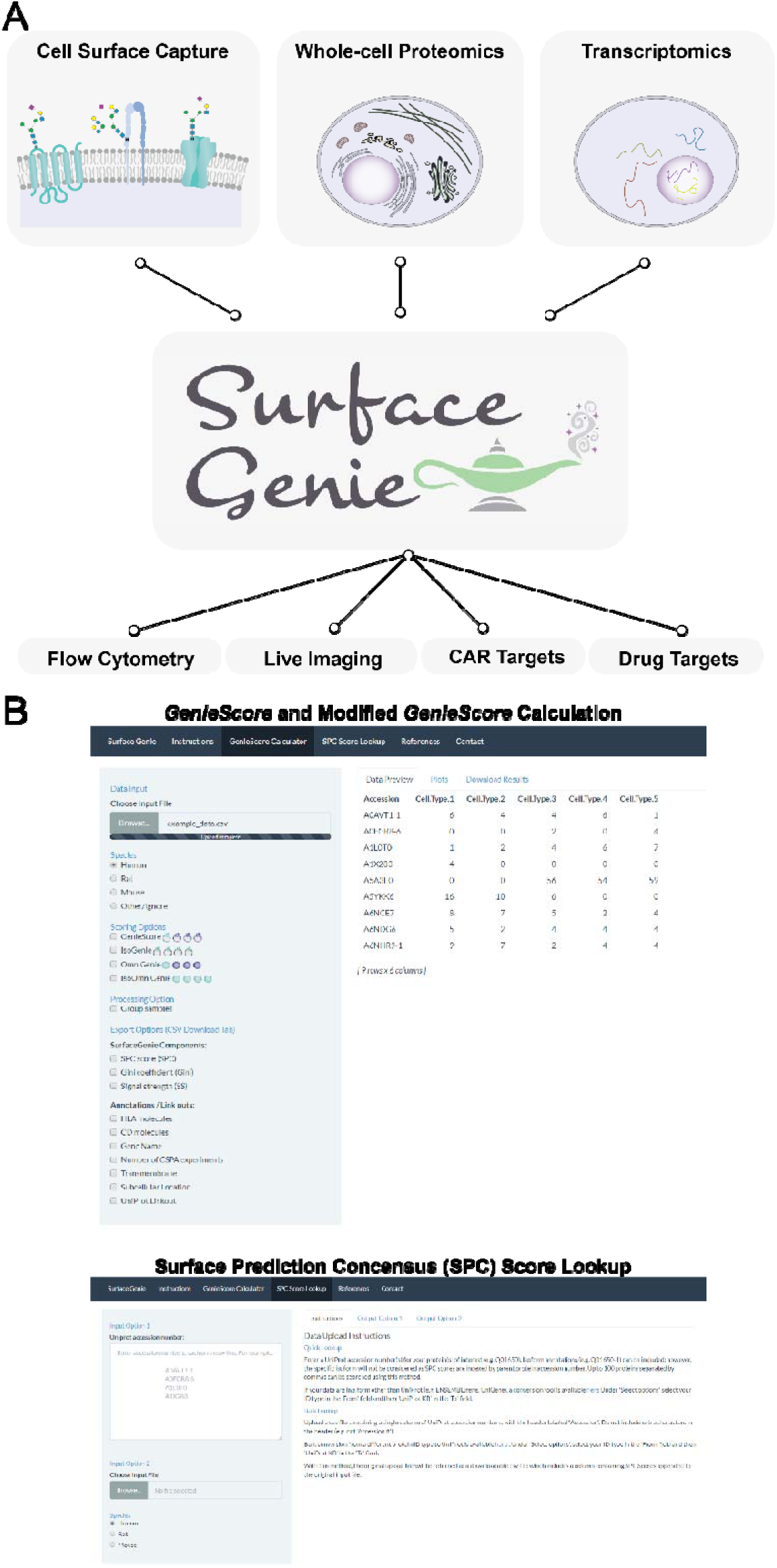
Overview of the utility of SurfaceGenie and screen captures from the web application. (A) A schematic depicting the tested inputs and potential applications of SurfaceGenie, a web-based application which calculates *GenieScore* permutations from user-input data. (B) Screen captures of the different modes of use for the SurfaceGenie web application.

## Discussion

Despite the central role cell surface proteins play in maintaining cellular structure and function, the cell surface is not well documented for most human cell types. There is currently no comprehensive reference repository of experimentally determined cell surface proteins cataloged by individual human cell types that can be used as a baseline for comparison to experimentally-perturbed or diseased phenotypes. Although specialized proteomic approaches allow for probing the occupancy of the cell surface, the sample requirements and technical sophistication often preclude widespread application, and quantitation is challenging. To overcome these challenges, predictions of surface localization can enable insights from more easily implemented proteomic and transcriptomic approaches, which can be performed on smaller sample sizes. However, with technologies that allow for ‘omic’ evaluation of individual cell types, there is a need to develop methodologies to prioritize the value contained within these studies in order to extract useful knowledge from acquired data.

Here, we describe the development of *GenieScore,* a prioritization approach that integrates a predictive metric regarding surface localization with experimental data to rank-order proteins which may be useful as cell surface markers. We demonstrate that *GenieScore* is compatible with quantitative data from CSC, WCL, and RNA-Seq experiments and is a useful strategy by which to integrate multiple sources of data for candidate marker prioritization. The design of *GenieScore* was intended for comparing data collected of various cell types or conditions within the same batch. Comparisons involving data from multiple sources may benefit from utilization of batch correction (Espin-Perez et al., 2018) or signal harmonization methods (Pino et al., 2018) for transcriptomic or proteomic data, respectively. SurfaceGenie, a web-based application, was developed to enable the calculation of *SPC scores* and *GenieScores*, and the various permutations thereof, from user-input data. SurfaceGenie also supplements the data with annotations relevant for marker selection.

Beyond immunophenotyping, SurfaceGenie is expected to facilitate the identification of valuable drug targets as the features of cell surface markers (*e.g.* surface localization and cell-type specificity) are also advantageous when designing efficient and specific therapies. Independent of *GenieScore*, the ability to query *SPC scores* within SurfaceGenie can deliver value in-and-of-itself, providing users with an additional resource to interrogate surface localization for proteins which are not yet characterized experimentally. However, whether an expressed protein is localized to the cell surface on a specific cell type in a specific experimental or biological condition remains difficult to predict. This is especially true for proteins that 1) lack traditional sequence motifs (*e.g.* signal peptides), 2) are only trafficked to the cell surface upon ligand binding (*e.g.* glucose transporter 3, GLUT3), or 3) have proteoforms that exhibit different subcellular localization than the canonical version of a protein for which predictions are typically based upon. For these reasons, experimental workflows that provide capabilities for discovery (*i.e.* not limited to available affinity reagents) while providing experimental evidence of cell surface localization on a particular cell type of interest with a specific context (*e.g.* experimental condition, disease state) will remain invaluable.

In conclusion, we anticipate that SurfaceGenie will enable effective prioritization of informative candidate cell surface markers to support a broad range of research questions, from mechanistic to disease-related studies. The candidates prioritized using SurfaceGenie are expected to be of use to a range of applications including immunophenotyping, immunotherapy, and drug targeting.

## Materials and Methods

All experimental details are provided in Supporting Information.

### Cell culture

Human lymphocyte cell lines (Ramos, HG-3, RCH-ACV, Jurkat) were cultured and passaged as previously described (Haverland et al., 2017). α TC1 clone 6 (CRL-2934; ATCC, Manassas, VA) and β-TC-6 (CRL-11506; ATCC) cells were maintained at 37°C and 5% CO_2_, cultured in Dulbecco’s Modified Eagle’s Medium supplemented with 10% heat-inactivated fetal bovine serum containing 16.6 mM or 5.5 mM glucose, respectively.

### Cell Lysis, Protein Digestion, and Peptide Cleanup

For WCL analysis of lymphocytes, cell pellets were lysed in 100 mM ammonium bicarbonate containing 20% acetonitrile and 40% Invitrosol (Thermo Fisher Scientific, Waltham, MA), digested with trypsin (Promega, Madison, WI) overnight, and cleaned by SP2 (Waas, Pereckas, Jones Lipinski, Ashwood, & Gundry, 2019). Peptides were quantified using Pierce Quantitative Fluorometric Peptide Assay (Thermo Fisher Scientific) according to manufacturer’s instructions on a Varioskan LUX Multimode Microplate Reader and SkanIt 5.0 software (Thermo Fisher Scientific). For CSC analysis of mouse islet cell lines, samples were prepared as previously described (Boheler et al., 2014; Gundry et al., 2012; Haverland et al., 2017).

### Mass Spectrometry Acquisition and Analysis

Lymphocyte peptides and CSC samples of mouse islet cell types were analyzed by LC-MS/MS using a Dionex UltiMate 3000 RSLCnano system (Thermo Fisher Scientific) in line with a Q Exactive (Thermo Fisher Scientific). Lymphocyte samples were prepared as 50 ng/µL total sample peptide concentration with Pierce Peptide Retention Time Calibration Mixture (PRTC, Thermo Fisher Scientific) spiked in at a final concentration of 2 fmol/µL and queued in blocked and randomized order with two technical replicates analyzed per sample. CSC samples of mouse islet cell types were analyzed as described (Mallanna, Cayo, Twaroski, Gundry, & Duncan, 2016; Mallanna, Waas, Duncan, & Gundry, 2016). MS data were analyzed using Proteome Discoverer 2.2 (Thermo Fisher Scientific) and SkylineDaily (v4.2.1.19095) (Schilling et al., 2012). One way ANOVA of CSC and WCL PSMs were performed using R (Team, 2018).

### Construction of a consensus dataset of predicted surface proteins

Four published surfaceome datasets (Bausch-Fluck et al., 2018; da Cunha et al., 2009; Diaz-Ramos et al., 2011; Town et al., 2016), each of which used a distinct methodology to bioinformatically predict the subset of the proteome which can be surface localized, were concatenated into a single consensus dataset. The ‘retrieve/mapping ID’ function within UniProt (www.uniprot.org) was used to convert the gene names provided in the published datasets to UniProt Accession numbers. Ambiguous matches were clarified by any supplementary information provided in the datasets in addition to gene name (*e.g.* alternate name, molecule name, chromosome).

### GenieScore – A mathematical representation of surface marker potential

An equation was developed to mathematically represent key features deemed relevant when considering whether a protein has high potential to qualify as a cell surface marker for distinguishing between cell types or experimental groups. The equation, which returns a metric termed *GenieScore*, is the product of 1) the *SPC scores* (described above); 2) *signal dispersion*, a measure of the disparity in observations among investigated samples that is mathematically equivalent to the square of the normalized Gini coefficient (Giugni, 1912); and 3) *signal strength*, a logarithmic transformation of the experimental data (*e.g.* number of PSMs, MS1 peak area, FKPM, or RKPM). A thorough definition and rationalization of the individual equation terms is provided in Supporting Information.

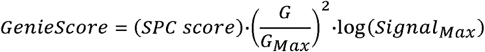

### Application of GenieScore

Details for the strategies applied to calculate *GenieScores* for each study are provided in Supporting Information.

### SurfaceGenie Web application

A web application for accessing SurfaceGenie was developed as an interactive Shiny app written in R and is available for use at www.cellsurfer.net/surfacegenie. Source code is available at www.github.com\GundryLab\SurfaceGenie.

## Supporting Information

1. Supplemental Methods
2. Figure S1 – Benchmarking of *Surface Prediction Consensus* (*SPC*) database against CSPA and HyperLOPIT annotations
3. Figure S2 – Visual depiction of Gini coefficient calculation and examples of *GenieScore* calculations
4. Figure S3 – Hierarchical clustering applied to all and predicted surface proteins for Cell Surface Capture and whole-cell lysate data
5. Figure S4 – Correlation of *GenieScore* experimental terms with statistical significance and correlation of *GenieScores* calculated using PSMs or MS1-based peak area
6. Dataset S1 – (1) Human SPC dataset, (2) Mouse SPC dataset, (3) Rat SPC dataset,
7. Dataset S2 - (1) Lymphocyte WCL data with *GenieScores*, (2) Lymphocyte CSC data with *GenieScores*, (3) *GenieScores* for proteins common to CSC and WCL
8. Dataset S3 – (1) CSC data on MCF10A KRAS^G12V^ and empty vector controls with *GenieScores*, (2) RNA-Seq data on MCF10A KRAS^G12V^ and empty vector controls with *GenieScores*
9. Dataset S4 – (1) Human dermal fibroblast and stem cell CSC data with *GenieScores*
10. Dataset S5 – (1) CSC data on mouse α and β cells with *GenieScores*, (2) RNA-Seq data on mouse α and β cells with *GenieScores*
11. Dataset S6 – (1) Single-cell RNA-Seq on human islet cells with *GenieScores* and modified *GenieScores*
12. SurfaceGenie User Guide – (1) SurfaceGenie Overview, (2) GenieScore Calculator – Basics and Tutorial, (3) SPC Score Lookup – Basics and Tutorial, (4)Additional Information

## Author Contributions

R.L.G. and M.W. conceived the study; R.L.G. supervised the study; M.W. developed the algorithms and designed and performed MS experiments; S.S. developed the R code; S.S. and J. L. developed the web application; R.A.J.L., P.A.H., J.A.C., performed analyses of mouse islet cell lines, M.W. and R.L.G. analyzed data; M.W. generated figures; M.W. and R.L.G. co-wrote the manuscript; All authors approved the final manuscript.

## Acknowledgements

This work was supported by the National Institutes of Health [R01-HL126785 and R01-HL134010 to R.L.G.; F31-HL140914 to M.W.; DK-052194 and AI-44458 to J.A.C.] and JDRF [2-SRA-2019-829-S-B to R.L.G. and J.A.C.]; S.S. is a member of the MCW-MSTP which is partially supported by a T32 grant from NIGMS, GM080202. Special thanks to Dr. Christopher Ashwood and Linda Berg Luecke for critical review of the manuscript and insightful discussions. Funding sources were not involved in study design, data collection, interpretation, analysis or publication.

