## Supplemental Data for "SurfaceGenie: a web-based application for prioritizing cell-type specific marker candidates"

**This PDF file includes:**

- Supplementary text
- Figs. S1 to S4
- Tables S1 – S2
- References for SI citations

**Other supplementary materials for this manuscript include the following:**

- Datasets S1-6
- SurfaceGenie User Guide

**Supplementary Information Text**

**Rationalization and description of *GenieScore* equation components.**

*Surface Protein Consensus (SPC) score* was generated from concatenating four individual human surfaceome databases and assigning a point for each of the individual datasets in which the protein was predicted to be localized to the cell surface. *SPC scores* range 0-4 such that proteins with more consensus of surface localization are prioritized over proteins with less consensus. Human, mouse and Rat SPC scores are in Dataset S1.

*Signal dispersion* is calculated for each protein based on the quantitative measurements from each cell type. First, the Gini coefficient, a measure of disparity, is calculated on the array of measurements. Next, this value is normalized by dividing by the maximum Gini coefficient possible, (1 – 1/N), where N is equal to the number of cell types. Finally, this value is squared to increase the weight assigned to this term. The values for this term range 0-1. Proteins with exactly equal measurements across cell types will score 0, proteins only observed in a single cell type will score 1. A visual depiction of Gini coefficient calculation is shown in Figure S2A. This measurement does not assume the normal distribution of data and requires no imputation of zero-values making it amenable to many types of quantitative measurements.

*Signal strength* is calculated for each protein based on the quantitative measurements from each cell type. First, the maximum measurement is calculated for each protein. Next, the $\log_{10}$ is calculated for 1 plus this value, in order to force all the values to be returned as positive numbers. This results in proteins at the lower limit of detection being of lower priority than those with a stronger signal, because it is expected that those of higher abundance will practically serve as more accessible markers for downstream technologies. *Signal strength* is not a bounded term and the range depends on the type of quantitative measurement.

**Modifications to the GenieScore equation.**

*IsoGenieScore* utilizes the same three calculations as *GenieScore* (see above), however, it uses (1 - *signal dispersion*). This prioritizes proteins with equal and intense measurements as opposed to those with disparate measurements.

*OmniGenieScore* is equal to the product of *signal dispersion* and *signal strength*. This prioritizes molecules with disparate measurements without considering the surface localization. As this score doesn't apply any protein-specific information, it can be calculated on any type on quantitative data.

*IsoOmniGenieScore* is equal to the product of (1 - *signal dispersion*) and *signal strength*. This prioritizes molecules with equal and intense measurements without considering the surface localization. As this score doesn't apply any protein-specific information, it can be calculated on any type on quantitative data.

**Methods**

*Cell lysis, protein digestion, and peptide cleanup*

For whole-cell lysate analysis of lymphocyte cell lines, pellets of 5 million cells were lysed in 500 µL of 2x Invitrosol (40% v/v; Thermo Fisher Scientific, Waltham, MA), 20% acetonitrile in 50 mM ammonium bicarbonate. Sample was sonicated (VialTweeter; Hielscher Ultrasonics, Teltow, Germany) by three ten-second pulses, set on ice for one minute, and then sonicated by three ten-second pulses. Samples were brought to 5mM TCEP and reduced for 30 min at 37°C on a Thermomixer at 1200 rpm. Samples were brought to 10 mM IAA and alkylated for 30 min at 37°C on a Thermomixer at 1200 rpm in the dark. 20 µg trypsin was added to each sample and was digested at 37°C overnight on a Thermomixer at 1200 rpm.

*Hierarchical clustering of lymphocyte whole-cell lysate (WCL) and Cell Surface Capture (CSC) data*

Data containing the number of peptide-spectrum matches were uploaded into SPSS (v. 22). Hierarchical clustering was performed using Phi-square measure of distance (appropriate for count data) and furthest neighbor (complete) linkage. Clustering was performed on entire dataset and then repeated for predicted surface proteins.

*MS1 Peak Area Quantification*

RAW and searched MS data for CSC and WCL were imported into SkylineDaily (v4.2.1.19095) (Schilling et al., 2012). For both CSC and WCL, peptide inclusion criteria were (1) fully tryptic, (2) no missed cleavages, (3) length 6-30, (4) exclude 25 N-terminal amino acids, and (5) no methionine residues. For WCL samples, all default, sequence-based exclusion criteria in Skyline except cysteine were further applied. For CSC, proteins with ≥ 3 peptides were selected for MS1-based quantification. For WCL, proteins with ≥ 5 peptides were candidates for MS1-based quantification. From among these candidates, the proteins with the top 15 and bottom 15 *GenieScores* were selected for MS1-based quantification.

*Lymphocyte whole-cell lysate (WCL) and Cell Surface Capture (CSC) data*:

WCL data were acquired and searched as part of this study using parameters in Tables S1 and S2. Searched WCL data were filtered to include only proteins with ≥2 unique peptides. RAW files were obtained from MassIVE (massive.ucsd.edu; accession number MSV000080532) for CSC experiments performed by Haverland *et al.* (Haverland et al., 2017) and re-searched using parameters in Table S2 (CSC Hi-Hi). Searched CSC data were filtered to include only proteins with ≥ 2 peptide-spectrum matches (PSMs) among all samples. All data are in Dataset S2.

*CSC and RNA-Seq data on MCF10A KRAS$^{G12V}$ and empty vector controls:*

CSC and RNA-Seq data were obtained from Supplemental Files 1 and 5, respectively, from Martinko *et al*. (Martinko et al., 2018). Only transcripts marked as “significantly different” were included in RNA-Seq analysis. As only $log_2$fold changes were provided for CSC data, these data were transformed to allow calculation of *signal dispersion*. The *signal strength* component calculated from FPKM values were used for both CSC and RNA-Seq analyses. All data are in Dataset S3.

*Human stem cell and dermal fibroblast CSC data*

RAW files were obtained from MassIVE (massive.ucsd.edu; accession number MSV000083846) for CSC experiments performed by Boheler *et al.* (Boheler et al., 2014). RAW files for embryonic stem cells (DR-11, DR-17, DR-27, DR-29), induced pluripotent stem cells (DR-28, DR-30, DR-31), and dermal fibroblasts (DR-12, RG-107, RG-108) were re-searched using parameters in Table S2 (CSC Hi-Lo). Searched CSC data were filtered to include only proteins with ≥ 2 PSMs among all samples. Embryonic and induced pluripotent stem cells were treated as a single group, averaging the number of PSMs for each protein. All data are in Dataset S4.

*α and β cell CSC and RNA-Seq data*

CSC data were acquired and RAW files were searched as part of this study according to parameters in Table S1 and S2 (CSC Hi-Hi). Searched CSC data were filtered to include only proteins occurring in ≥ 2 biological replicates. RNA-Seq data were obtained from Supplemental File 12 from Benner *et al.*(Benner et al., 2014). The combined *GenieScore* calculations were performed by first normalizing the PSMs and RPKM measurements individually to the maximum value for each protein. Next, the average of the normalized values was calculated for both α and β cells. Finally, the *signal dispersion* was calculated using these averaged, normalized measurements. The sum of the CSC and RNA-Seq *signal strength* values was used for the calculation of the combined *GenieScores*. All data are in Dataset S5.

*Islet cell single-cell RNA-Seq*

RNA-Seq data were obtained from Supplemental File 6 from Lawlor *et al*. (Lawlor et al., 2017). All data are in Dataset S6.

**Fig. S1. Benchmarking *Surface Prediction Consensus (SPC)* scores**. (A-B) The distribution of confidence assignments within the Cell Surface Protein Atlas (CSPA) (Bausch-Fluck et al., 2015) across different *SPC scores* for human and mouse datasets depicted as bubble charts, where the size of the bubble represents the number of proteins in the intersection between the particular *SPC score* and CSPA annotations. (C) The distribution of annotations assigned by application of HyperLOPIT (Christoforou et al., 2016) to mouse stem cells across different *SPC scores* depicted as a bubble chart, where the size of the bubble represents the number of proteins in the intersection between the particular *SPC score* and HyperLOPIT annotation.

A

B

C

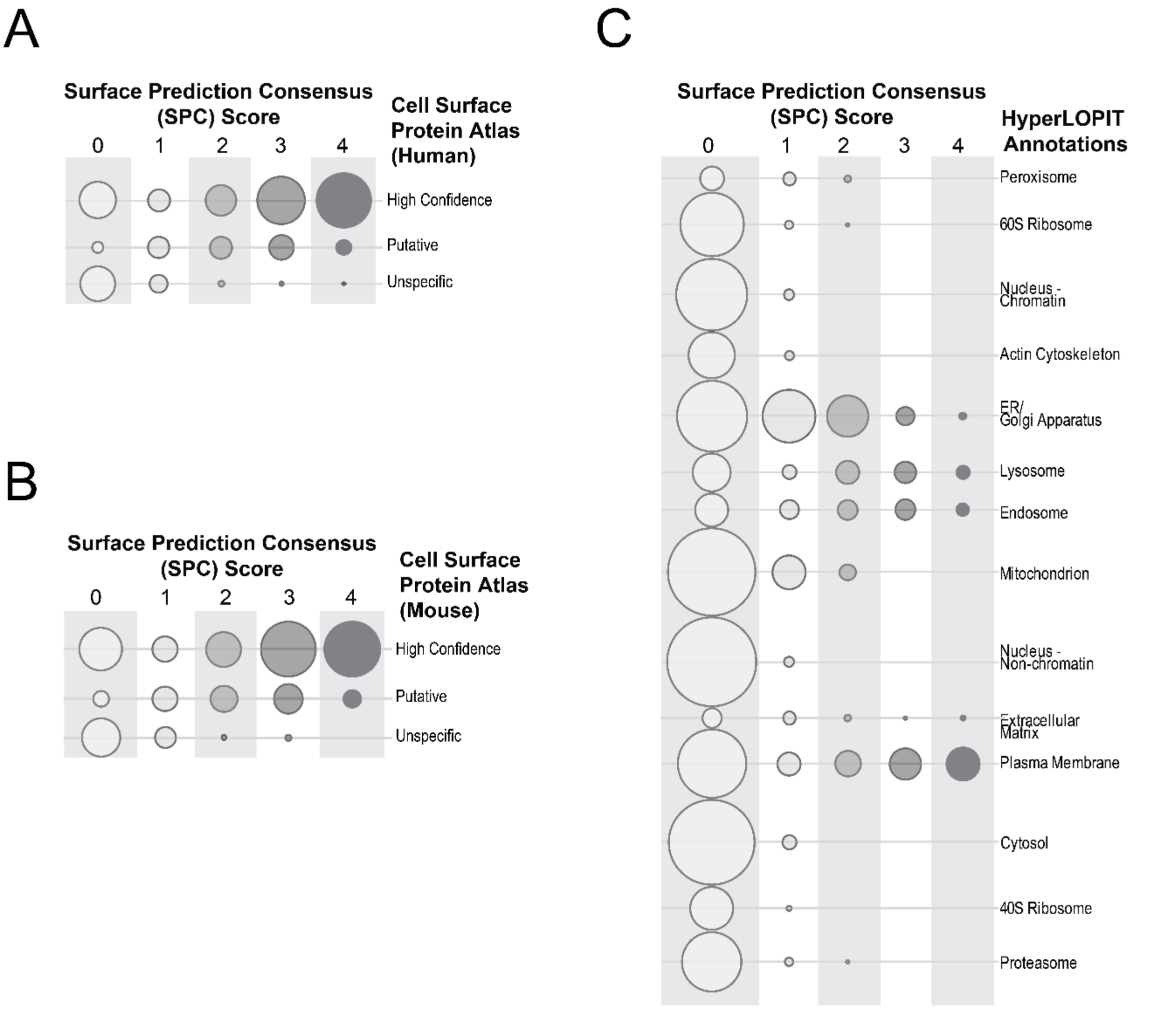

**Fig. S2. Gini coefficient calculation and examples of *GenieScore* calculations.** (A) The calculation of Gini coefficient is represented visually for two example proteins, where the gray shaded area represents the calculated disparity between measurements. Protein A has equal measurements in each sample type. Tracing the addition to cumulative signal from each sample results in the identity function '*y* = *x*' (*i.e.* each sample/measurement pair contributes equally). The contributions to total Protein B signal are disparate among the cell types, meaning that sample/measurement pair contributes unequally to their respective summed totals. The gray shaded area for Protein B represents the discrete integral calculated from the identity function '*y* = *x*' to a point-to-point fit of the contributions to cumulative signal. The gray shaded area is used to calculate the Gini coefficient. As Protein A has an area of 0, it also has a Gini coefficient of 0. (B) Examples of *GenieScores* are shown for three pairs of proteins, which differ with respect to one of the individual components of the *GenieScore* equation.

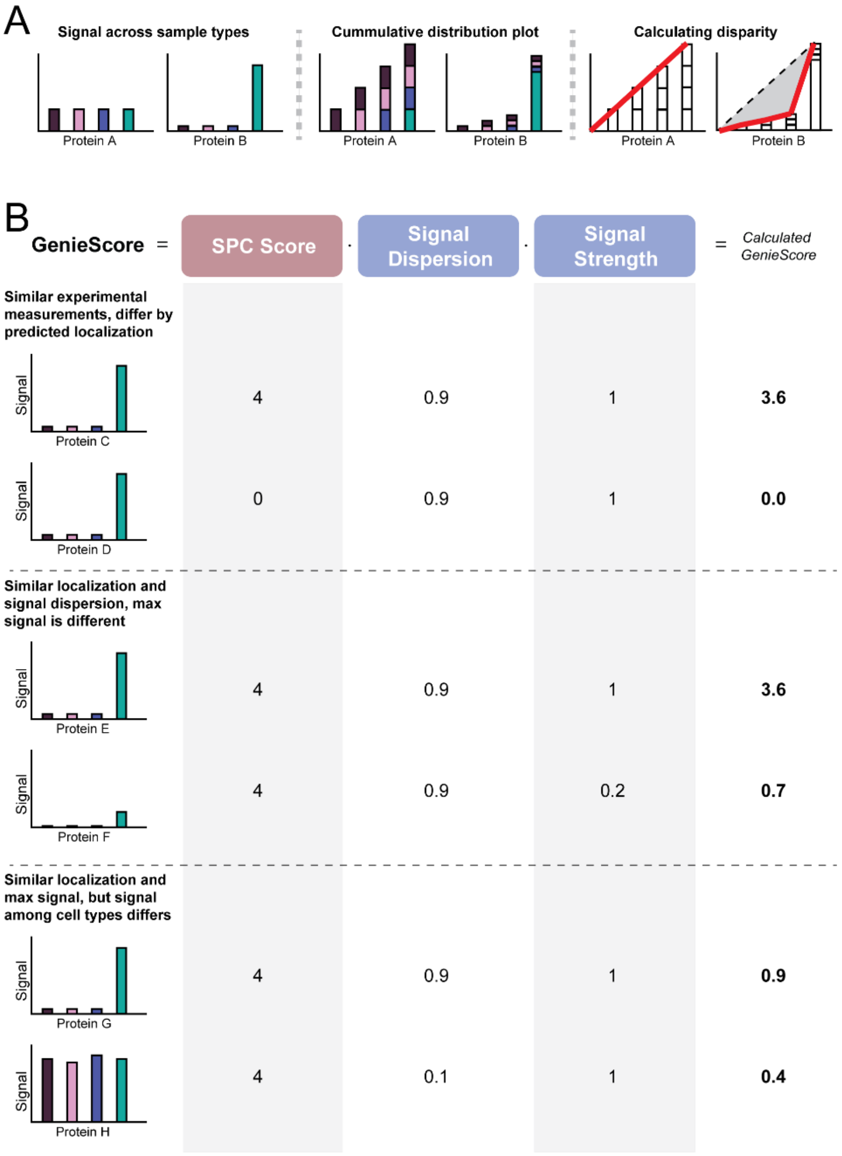

**Fig. S3. Hierarchical clustering of lymphocyte Cell Surface Capture and Whole-cell Lysate data.** Dendrograms depicting the relationships inferred by hierarchical clustering. All three biological replicates cluster for each of the four lymphocyte cells lines whether using all identified proteins or the subset of proteins predicted to be surface-localized by *SPC scores*.

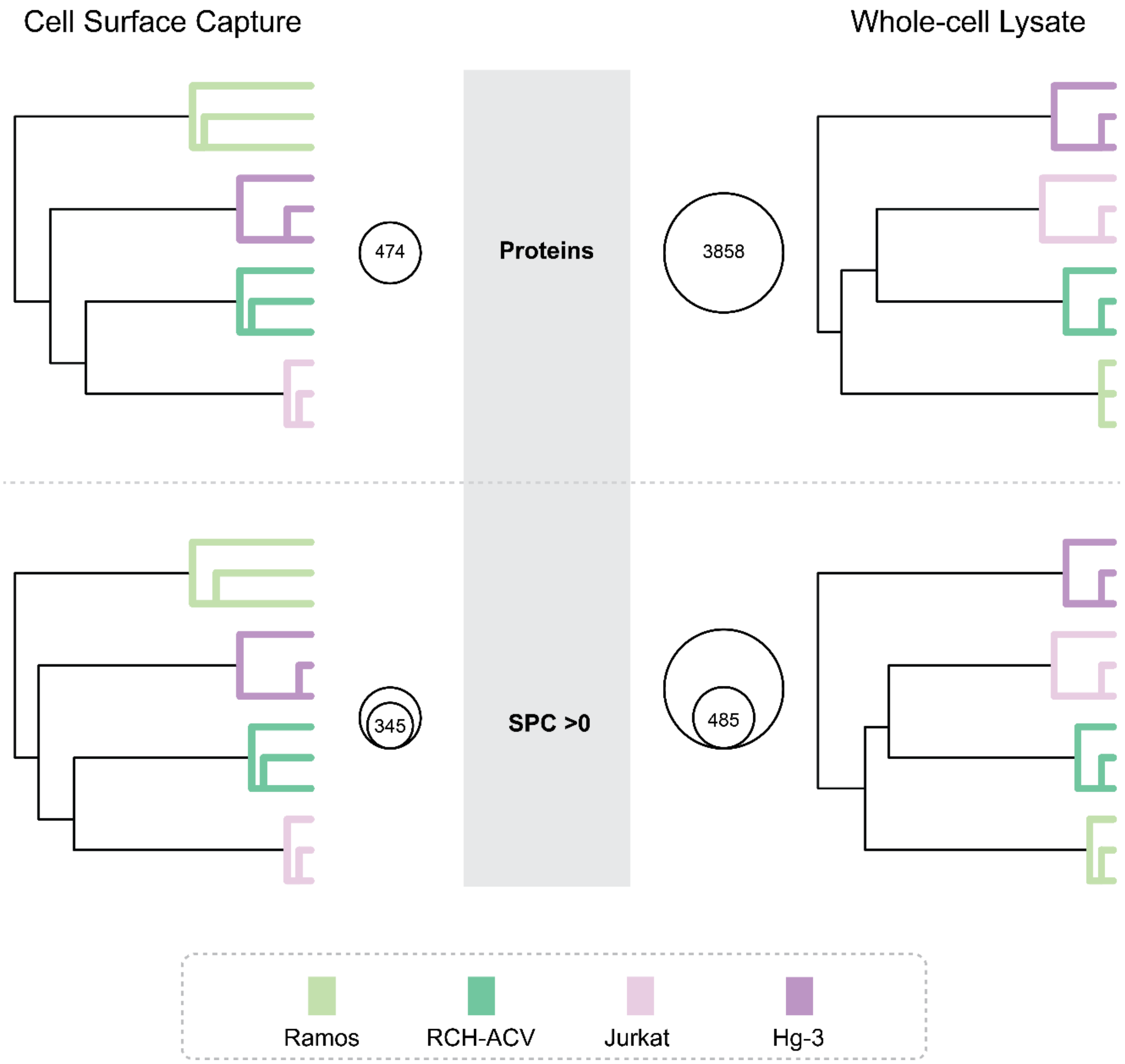

**Fig. S4. Correlations of *GenieScore*s and *GenieScore* components.** *GenieScores* calculated MS1-based peak area plotted against *GenieScores* for the same proteins calculated using peptide-spectrum matches shown with calculated Spearman's correlation for (A) whole-cell lysate (WCL) and (B) Cell Surface Capture (CSC) data. The product of signal distribution and signal strength plotted against the statistical significance calculated using a one-way ANOVA shown with calculated Spearman's correlation for (C) WCL and (D) CSC data.

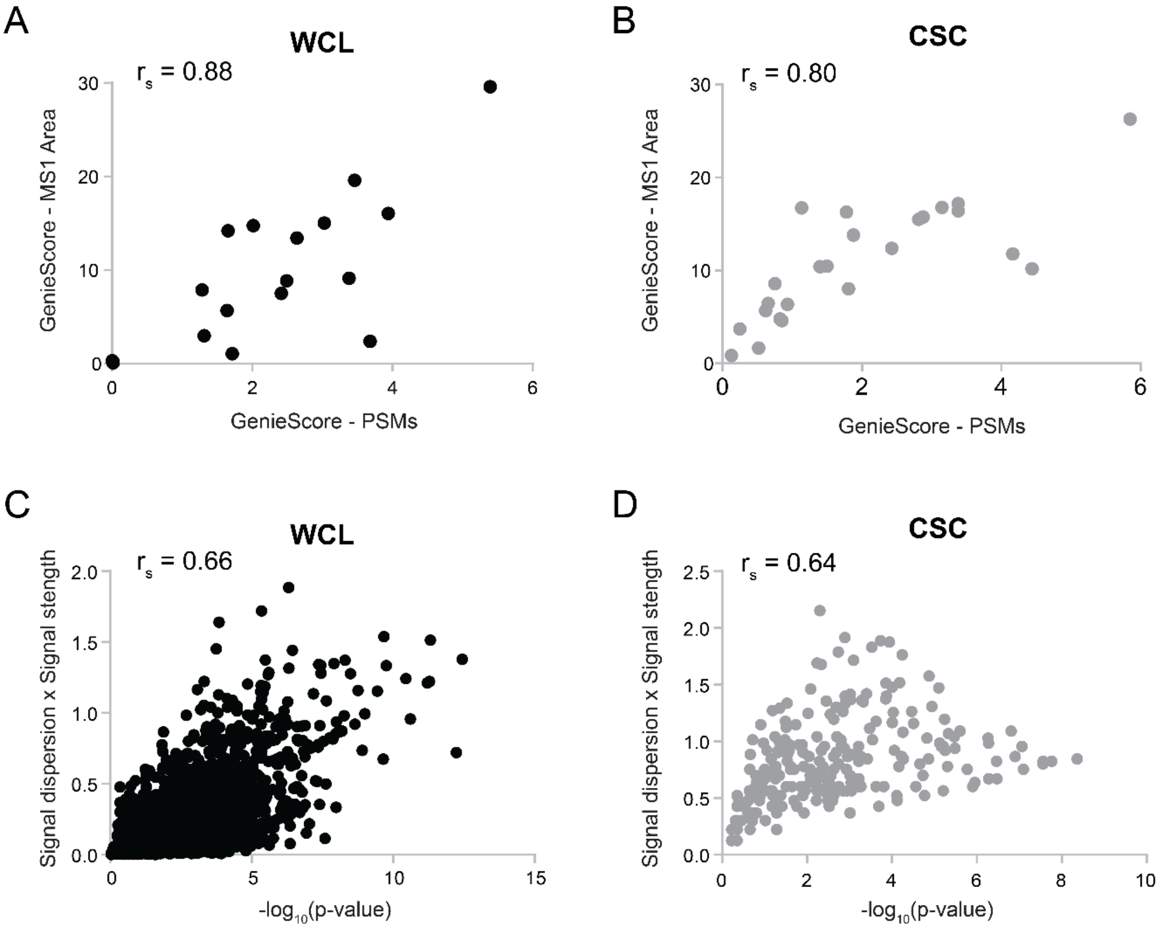

**Table S1. Mass spectrometry acquisition settings**

| | Whole-cell Lysate | Cell Surface Capture |
|---|---|---|
| ***Injection Mode*** | Full Loop | uL PickUp |
| ***Sample Loop*** | 20 µL | 20 µL |
| ***Stationary Phase*** | Acclaim PepMap C 18 100 Å, 75 µm, 2 µm, 25 cm | Michrom Bioresources Magic C18AQ 200 Å, 3 µm, 10 cm |
| ***LC Solvent A*** | 100% $H_2O$,<br>0.1% formic acid | 100% $H_2O$,<br>0.1% formic acid |
| ***LC Solvent B*** | 80% MeCN,<br>0.1% formic acid | 80% MeCN,<br>0.1% formic acid |
| ***LC Gradient*** | 7-7% B in 5 min<br>7-28% B in 123 min<br>28-40% B in 25 min<br>40-98% in 3 min | 2-2% B in 10 min<br>2-35% B in 40 min<br>35-98% B in 10 min |
| ***LC Flow Rate*** | 300 nL/min | 300 nL/min |
| ***Mass Spectrometer*** | Thermo Orbitrap Q Exactive | Thermo Orbitrap Q Exactive |
| ***Method Type*** | Data dependent MS2, Top15 | Data dependent MS2, Top15 |
| ***Spray Voltage*** | 2 kV | 3.8 kV |
| ***$MS^1$ Detector*** | Orbitrap | Orbitrap |
| ***$MS^1$ scan range*** | 350-1600 m/z | 300-1600 m/z |
| ***$MS^1$ resolution*** | 70,000 @ 200 m/z | 70,000 @ 200 *m/z* |
| ***$MS^1$ AGC Target*** | 1e6 | 1e6 |
| ***$MS^1$ Maximum IT*** | 50 ms | 50 ms |
| ***$MS^2$ Detector*** | Orbitrap | Orbitrap |
| ***$MS^2$ resolution*** | 17,500 @ 200 m/z | 17,500 @ 200 m/z |
| ***Isolation Window*** | 2.0 m/z | 2.0 m/z |
| ***$MS^2$ AGC Target*** | 5e4 | 1e5 |
| ***$MS^2$ Maximum IT*** | 110 ms | 110 ms |
| ***Activation Type / Collision Energy*** | HCD 27% | HCD 27% |
| ***Minimum AGC Target.*** | 5.0e2 | 1.0e3 |
| ***Intensity Threshold*** | 4.5e3 | 9.1e3 |
| ***Dynamic Exclusion*** | 30 s | 60 s |

### Table S2. Peptide search and post-search validation parameters

| ***Sample*** | **Whole-cell lysate** | **Cell Surface Capture (Hi-Hi)** | **Cell Surface Capture (Hi-Lo)** |
|---|---|---|---|
| ***Platform*** | ProteomeDiscoverer 2.2 | ProteomeDiscoverer 2.2 | ProteomeDiscoverer 2.2 |
| ***Search Algorithm*** | SequestHT | SequestHT | SequestHT |
| ***Validation*** | Percolator<br>Peptide Validator<br>Protein FDR Validator | Percolator<br>Peptide Validator<br>Protein FDR Validator | Percolator<br>Peptide Validator<br>Protein FDR Validator |
| ***Database*** | SwissProt; Human; created 6/7/2017 | SwissProt; Human; created 6/7/2017 or SwissProt; Mouse; created 11/30/2018 | SwissProt; Human; created 6/7/2017 |
| ***Enzyme (semi/full)*** | Trypsin (full) | Trypsin (semi) | Trypsin (semi) |
| ***Missed Cleavages*** | 2 | 2 | 2 |
| ***Precursor mass tolerance*** | 10 ppm | 10 ppm | 10 ppm |
| ***Fragment mass tolerance*** | 0.02 Da | 0.02 Da | 0.6 Da |
| ***Static Modifications*** | Carbamidomethyl (C) | Carbamidomethyl (C) | Carbamidomethyl (C) |
| ***Dynamic Modifications*** | Oxidation (M),<br>Acetylation (N-term) | Oxidation (M),<br>Acetylation (N-term)<br>Deamidation (N) | Oxidation (M),<br>Acetylation (N-term)<br>Deamidation (N) |
| ***Target FDR (Strict):*** | 0.01 | 0.01 | 0.01 |
| ***Target FDR (Relaxed):*** | 0.05 | 0.05 | 0.05 |
| ***Validation basis*** | q-Value | q-Value | q-Value |

**Additional dataset S1 (separate file – descriptions of separate tabs are below)**

(1) Human SPC dataset
(2) Mouse SPC dataset
(3) Rat SPC dataset

**Additional dataset S2 (separate file – descriptions of separate tabs are below)**

(1) Lymphocyte WCL data with *GenieScores*
(2) Lymphocyte CSC data with *GenieScores*
(3) *GenieScores* for proteins common to CSC and WCL

**Additional dataset S3 (separate file – descriptions of separate tabs are below)**

(1) CSC data on MCF10A $KRAS^{G12V}$ and empty vector controls with *GenieScores*
(2) RNA-Seq data on MCF10A $KRAS^{G12V}$ and empty vector controls with *GenieScores*

**Additional dataset S4 (separate file – descriptions of separate tabs are below)**

(1) Human dermal fibroblast and stem cell CSC data with *GenieScores*

**Additional dataset S5 (separate file – descriptions of separate tabs are below)**

(1) CSC data on mouse α and β cells with *GenieScores*
(2) RNA-Seq data on mouse α and β cells with *GenieScores*

**Additional dataset S6 (separate file – descriptions of separate tabs are below)**

(1) Single-cell RNA-Seq on human islet cells with *GenieScores* and modified *GenieScores*

**SurfaceGenie User Guide (separate file – descriptions of sections are below)**

(1) SurfaceGenie Overview
(2) GenieScore Calculator – Basics and Tutorial
(3) SPC Score Lookup – Basics and Tutorial
(4) Additional Information
